# Behavioral signatures of Y-like neuronal responses in human vision

**DOI:** 10.1101/2022.02.14.480142

**Authors:** Ana L. Ramirez, Lowell W. Thompson, Ari Rosenberg, Curtis L. Baker

## Abstract

Retinal ganglion cells initiating the magnocellular/Y-cell visual pathways respond nonlinearly to high spatial frequencies (SFs) and temporal frequencies (TFs). This nonlinearity is implicated in the processing of contrast modulation (CM) stimuli in cats and monkeys, but its contribution to human visual perception is not well understood. Here, we evaluate human psychophysical performance for CM stimuli, consisting of a high SF grating carrier whose contrast is modulated by a low SF sinewave envelope. Subjects reported the direction of motion of CM envelopes or luminance modulation (LM) gratings at different eccentricities. The performance on SF (for LMs) or carrier SF (for CMs) was measured for different TFs (LMs) or carrier TFs (CMs). The best performance for LMs was at lower TFs and SFs, decreasing systematically with eccentricity. However, performance with CMs was bandpass with carrier SF, largely independent of carrier TF, and at the highest carrier TF (20 Hz) decreased minimally with eccentricity. Since the nonlinear subunits of Y-cells respond better at higher TFs compared to the linear response components and respond best at higher SFs that are relatively independent of eccentricity, these results suggest that behavioral tasks employing CM stimuli might reveal nonlinear contributions of retinal Y-like cells to human perception.

## Introduction

Information about the visual world is provided to the brain through the responses of retinal ganglion cells (RGCs). There are many categorically distinct types of RGCs^1^, and establishing their functional contributions to perception has been challenging. Early studies in the cat characterized two principal types of RGCs, X-beta cells and Y-alpha cells^2-4^, which comprise key elements along the retino-geniculate pathway. Later studies confirmed that X-beta cells and Y-alpha cells were a common feature in all mammals examined so far^5^. In the primate, two main types of RGCs, midget and parasol cells, which are the origin of the parvocellular (P) and magnocellular (M) geniculocortical pathways were well characterized. Similar to primate midget cells, X-beta cells are compact neurons with small axons and dendritic fields. Primate parasol cells are like Y-alpha cells in having large receptive and dendritic fields. However, it was not immediately clear whether parasol cells possessed the characteristic frequency-doubling nonlinearity of Y-alpha cells, and it was therefore ambiguous whether primates possess a RGC counterpart of cat Y-alpha cells. Later research demonstrated that parasol and smooth/upsilon cells, which we collectively refer to here as Y-like cells, exhibit both anatomical and functional properties of cat Y-alpha cells including nonlinear responses to visual stimuli^6-8^. The functional role of Y-like cells’ nonlinear responses is not well understood in human vision.

Classic experiments showed that selective lesions of the magnocellular pathway decrease contrast sensitivity for visual stimuli at high temporal frequencies (TFs) and low spatial frequencies (SFs) but do not affect color vision^9-11^. However, those experiments did not investigate the contributions of the nonlinear responses of Y-like cells. Y-like cells are distinctive in that, at low SFs, they show a linear first Fourier harmonic (F1) response at the temporal frequency of a drifting grating, but when contrast-reversing gratings at high SFs are presented, they display a prominent second Fourier harmonic (F2) nonlinear response. This pattern of SF-dependence of F1 and F2 responses, often referred to as the “Y-cell signature", is characteristic of the nonlinear summation of Y-cells^4^. The F1 response is dependent on the spatial phase of a contrast-reversing grating, whereas the F2 nonlinear response is phase-independent. This characteristic phase-invariance of the F2 response of Y-like cells reflects the activation of nonlinear subunits originating from a specialized class of bipolar cells^7,8,12^. In neurophysiology experiments in primate retina, the estimated F1 linear center receptive fields clearly decrease systematically with eccentricity, as might be expected. In contrast, the F2 nonlinear center receptive fields maintain a relatively similar size across a large range of retinal eccentricities^8^, although the differences have not been compared quantitatively.

Contrast modulation (CM) stimuli generally consist of a carrier sine wave grating at a high SF, whose contrast is modulated by a sinewave envelope at a low SF. Neurophysiology studies have demonstrated that cortical neurons responding to CM stimuli exhibit selectivity for the carrier that is similar to the selectivity of nonlinear subunits of subcortical Y-cells^13-15^. These results support the idea that cortical CM responses are critically dependent on Y-cell inputs^14,16,17^.

Here we leveraged the Y-like carrier response properties of cortical neurons to CM stimuli to create a novel approach capable of revealing the nonlinear contributions of Y-like cells to human perception. We employed CM stimuli with a carrier at high spatial and temporal frequencies and an envelope at low spatial and temporal frequencies. Judgments of the direction of motion for conventional luminance modulation (LM) gratings were consistent with conventionally described linear mechanisms, whereas the results for envelope (CM) motion were consistent with Y-like nonlinear processing.

## Materials and Methods

### Subjects

Six healthy subjects (aged 23 to 35 years; 3 males, 3 females) participated in the study. Five were naive to the aims of the study (students from McGill University) and one was an author (ARH). All subjects had a monocular visual acuity or best-corrected acuity of at least 20/25 and reported no history of ophthalmological diseases or surgeries. All procedures were approved by the Research Ethics Board of the Research Institute of McGill University Health Center, and were performed according to relevant guidelines and regulations. All subjects gave written informed consent to participate.

### Apparatus

Visual stimuli were produced on a Macintosh computer (MacPro 4.1, MacOS X 10.6.8, 2×2.8 GHz Quad-Core, 24 GB RAM) with custom software written in MATLAB (MathWorks, Inc.) using Psychophysics Toolbox, version 3.0.10^18-20^. The stimuli were presented on a cathode-ray tube (CRT) monitor (Iiyama, 39.5 × 29.5 cm, 120 Hz, 1024 × 768 pixels) at a viewing distance of 221 cm. To achieve linearization and high monochromatic luminance level resolution, we first measured the CRT gamma nonlinearity with a photometer (United Detector Technology S370), and then used the LOBES Video Switcher^21^ that combines blue and attenuated red outputs from the graphics card to generate 16 bits of voltage resolution.

### Stimuli

The visual stimuli were presented at the center of the monitor’s screen within a cosine-tapered circular window of 3.5° of visual angle, on a uniform background at the mean luminance of the stimulus. We presented both first-order, luminance modulation (LM) gratings as well as second-order, contrast modulation (CM) gratings (**Supplementary Video S1**). A drifting LM grating was defined by:

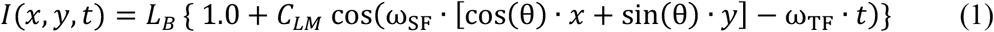

where *I*(*x, y, t*)is the luminance intensity of a pixel at spatial location (*x, y*,)at time *t, L*_*B*_ is the background (and mean) luminance, *C*_*LM*_ is the Michelson contrast of the luminance modulation, *ω*_SF_ is the spatial frequency, *θ* is the orientation, and *ω*_TF_ is the temporal frequency. The sign of *ω*_TF_ determines the direction of motion. The orientation was always vertical, so the motion was either leftwards or rightwards (as in **Supplementary Video S1**).

A CM grating was defined by the contrast modulation of a high spatial frequency carrier grating by a low spatial frequency envelope grating (**Fig. 1**):

**Figure 1.**
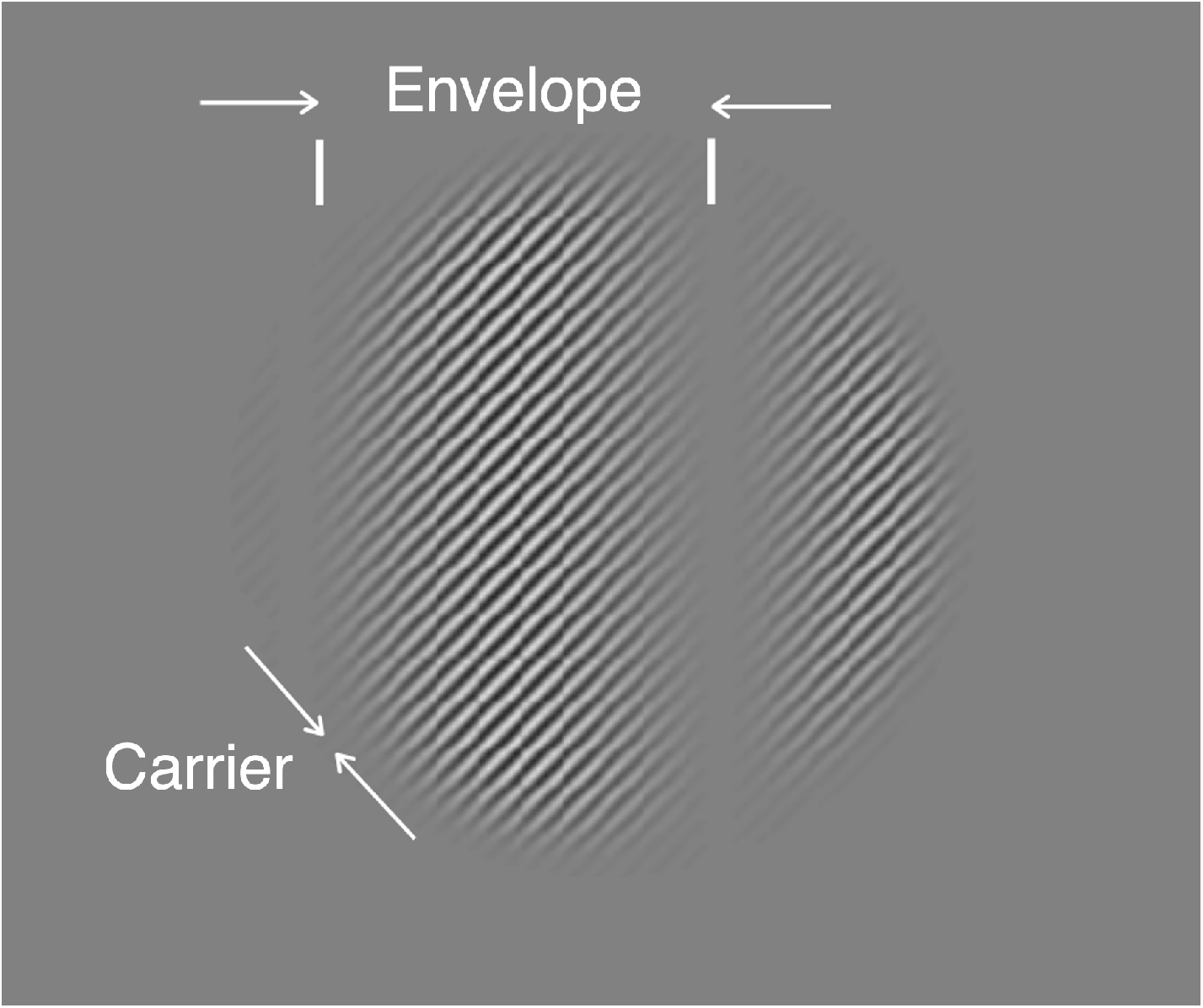
Example contrast modulation (CM) stimulus. The stimulus, presented within a cosine-tapered circular window of 3.5° of visual angle, consists of a high spatial frequency contrast-reversing carrier grating (right-oblique) whose contrast is modulated by a low spatial frequency, vertical sinewave envelope.

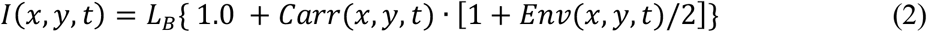

Here, *Carr*(*x, y, t*)is the carrier, which is a contrast-reversing grating with an orientation, spatial frequency, and temporal frequency that are defined independently of the envelope:

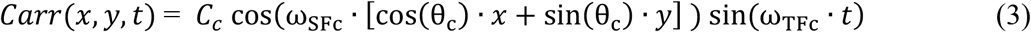

where *ω*_SFc_ is the carrier SF, *ω*_TFc_ is the carrier TF, θ_c_ is the carrier orientation and *C*_c_ is the carrier contrast. The carrier orientation was always right-oblique, as in **Figure 1**.

Likewise, *Env*(*x, y, t*)is the envelope, which is a drifting grating with an orientation, spatial frequency, and temporal frequency (direction) that are defined independently of the carrier:

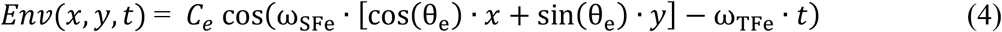

where *ω*_SFe_ is the envelope SF, *ω*_TFe_ is the envelope TF, θ_e_ is the envelope orientation and *C*_*e*_ is the envelope contrast, which henceforth we will refer to as the “modulation depth". The envelope orientation was always vertical, so the motion was either leftwards or rightwards (as in **Supplementary Video S1**).

A grey cardboard surround, 81 × 102 cm (width x height), framed the monitor and approximately matched the mean luminance (*L*_*B*_) of the stimuli on the CRT.

### Design and experimental procedures

Subjects were instructed to look at a fixation target which was placed in the upper right quadrant of the screen (**Fig. 2a,d**), such that the stimulus would be presented at 4.3° of eccentricity in the subject’s visual field, in the first experiment, and at varying eccentricities in the second experiment (see below). The fixation target and all visual stimuli were presented monocularly to the right eye for all subjects. On each trial, a stimulus was presented for 250 ms. The subject indicated the perceived direction of motion of the LM grating or CM envelope (**Fig. 2b,e**) by pressing a key, with no time limit, and with subsequent feedback (visual icon on the display screen) for incorrect responses. Performance was measured by the method of constant stimuli with seven logarithmically spaced values of spatial frequency (1.4, 2, 2.9, 4.2, 5.9, 8.4, and 12.5 cycles per degree (cpd)) and four values of temporal frequency (5, 10, 15, and 20 Hz). All experiments had a fixed contrast (*C*_*LM*_ and *C*_*c*_ from Eq 1 and 3, respectively) of 5% for LM gratings and 80% for CM envelope and carrier components, chosen based on pilot results to avoid ceiling effects in psychophysical performance and to compensate for the very different contrast sensitivities for the two kinds of stimuli^22,23^. A minimum of three blocks of 140 trials (20 trials per condition) was tested for each condition to provide a total of at least 60 trials per condition. Subjects generally completed one block in 5 to 10 minutes. In a second experiment, stimuli were presented at a fixed temporal frequency (20 Hz), at 2.1°, 4.3°, 6.4°, or 8.5° of eccentricity from the fixation target (**Fig. 2a,d**).

**Figure 2.**
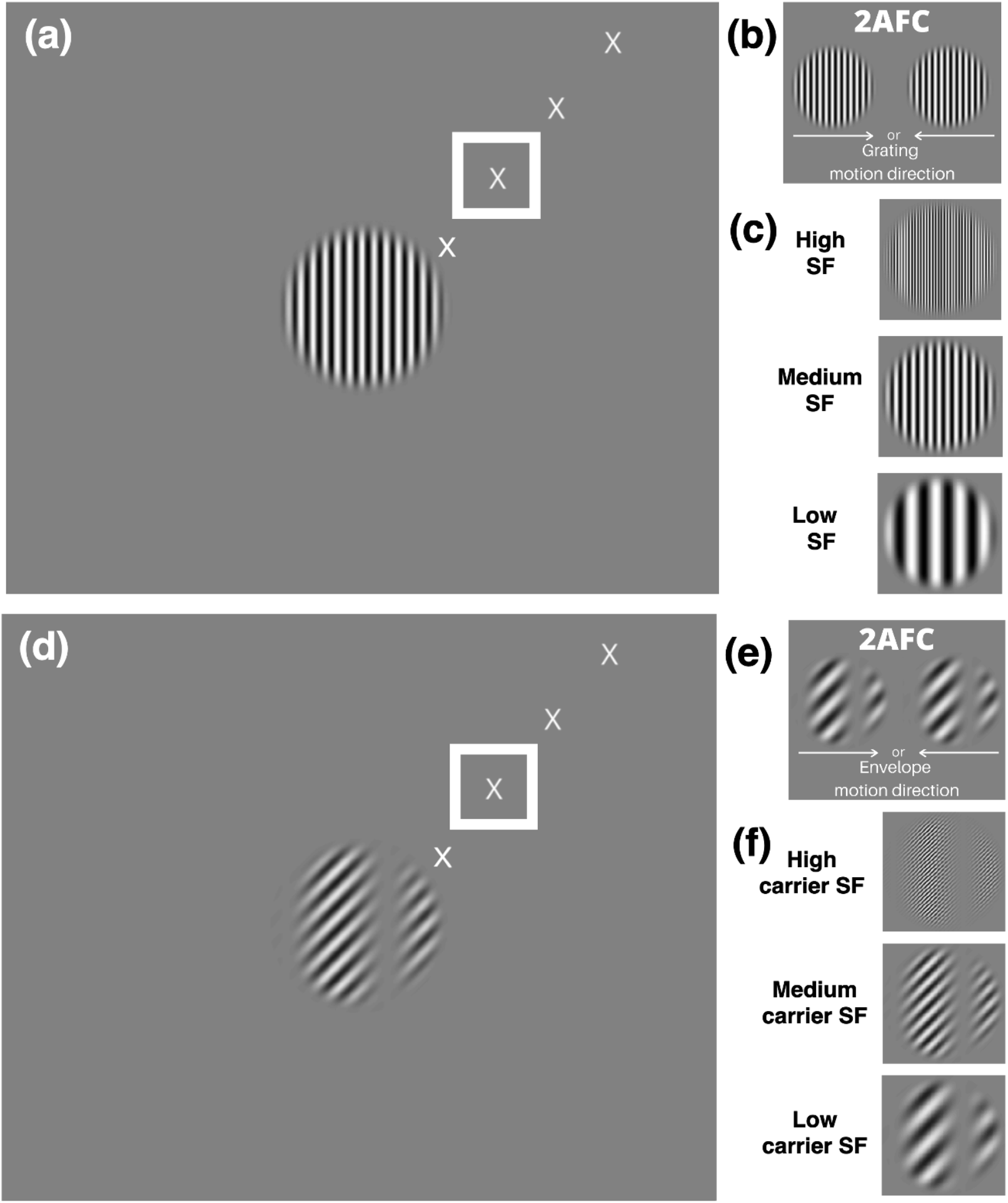
Behavioral task. (**a**) Example luminance modulation (LM) grating stimulus at the center of the screen, with different fixation targets (only one fixation target was visible on any given trial) that would place the stimulus at 2.1°, 4.3° (in a white square), 6.4°, and 8.5° of eccentricity. **(b)** Depiction of the 2-alternative forced choice (2AFC) task in which the subject reported the direction of motion (left vs right) of an LM grating at different spatial frequencies (SFs). **(c)**. Example LM stimuli at different SFs. **(d)** Example contrast modulation (CM) stimulus at the center of the screen, with the same fixation targets as in **a. (e)** Depiction of the 2AFC task in which the subject reported the direction of motion (left vs right) of a CM envelope at different carrier SFs. **(f)** Example CM stimuli at different carrier SFs. Fixation targets are slightly enlarged here for illustration purposes.

### Experimental Design and Statistical Analysis

Performance was determined by calculating each subject’s percent correct responses for each unique experimental condition. Standard errors were calculated for each condition based on the variance of the proportion in a binomial distribution resulting from Bernoulli trials: var(p) = p*(1-p)/N, where p is the proportion correct and N is the number of trials. For mean responses from all subjects (N=6), standard error of the mean was calculated. We used 2-way ANOVAs to test whether direction discrimination performance was significantly impacted by the stimulus SF, TF (or eccentricity), and their interaction.

To determine the SF acuity, i.e., the highest SF supporting a criterion performance threshold, for each subject and experimental condition, plots of percent correct data were fit with a logistic function. To do so, only the last 5 values of the high-SF fall-off were included (2.9, 4.2, 5.9, 8.4, and 12.5 cpd) to provide a monotonically decreasing function. The logistic curve-fitting was performed using Psignifit 4^24^. Threshold values of maximal SF (for LM) or carrier SF (for CM) were taken at the 75% correct level. Thresholds were included in an analysis if the 95% confidence interval did not include 0 (Experiment 1: 45/48 cases; Experiment 2: 41/48 cases). We used linear regression models to test whether the log SF acuity varied systematically as a function of TF (or eccentricity), stimulus type (LM or CM), and their interaction. We report the corresponding fixed-effect coefficients (b-values), F statistics, and p-values.

## Results

### Experiment 1: Spatial frequency dependence at different temporal frequencies

We hypothesized that direction discrimination performance for CM stimuli would remain relatively similar across a broad range of carrier TFs if the envelope motion was computed from the output of nonlinear Y-like cells rather than linear mechanisms. To test this prediction, we assessed performance across a variety of carrier SFs and TFs at a fixed eccentricity. To provide a comparative context for contrasting the results with those reflecting linear processing, we made similar measurements for luminance gratings (LM stimuli) over the same ranges of SFs and TFs, and at the same eccentricity. In particular, direction discrimination performance for LM stimuli was predicted to decrease systematically as a function of TF, as expected if perception were mediated by linear mechanisms generally thought to underlie performance for moving gratings^25-27^.

In each stimulus presentation of 250 ms duration, an LM or CM stimulus was presented in the center of the screen while the subject fixated a target 4.3° away from the stimulus (**Fig. 2a,d**, white square). The task was to report the perceived direction of motion of the LM grating or of the CM envelope. Within each block of trials, the LM grating SF or CM carrier SF was varied using the method of constant stimuli at a fixed TF (LM gratings) or carrier TF (CM envelopes). In separate blocks of trials, the dependence of performance on SF was measured for different values of TF (LM gratings) or carrier TF (CM envelopes).

We first analyzed performance with the LM stimuli (vertical gratings, drifting leftwards or rightwards), which were presented with a Michelson contrast of 5%. The percent correct performance as a function of SF is plotted for one subject in **Figure 3a** for a series of values of TF. The ability to correctly report the direction of motion at high SFs (2.9 to 5.9 cpd) was only possible at low TFs (5 to 10 Hz). Note that the performance fall-off with increasing SF was faster at successively higher TFs. **Figure 3b-f** plots the results for 5 additional subjects, who showed similar patterns. The average performance across all 6 subjects is plotted in **Figure 3g**. In general, performance fell off with increasing SF, and was more rapid for higher TF values (except at the two lowest TF values, at which performance was similar). These observations were confirmed with a 2-way ANOVA (**Fig. 3g**), which showed a significant main effect of SF, F(6,168) = 117.02, p = 2.39×10^−57^; a significant main effect of TF, F(3,168) = 20.08, p = 3.59×10^−11^; and a significant interaction effect, F(18,168) = 3.3, p = 2.43×10^−5^.

**Figure 3.**
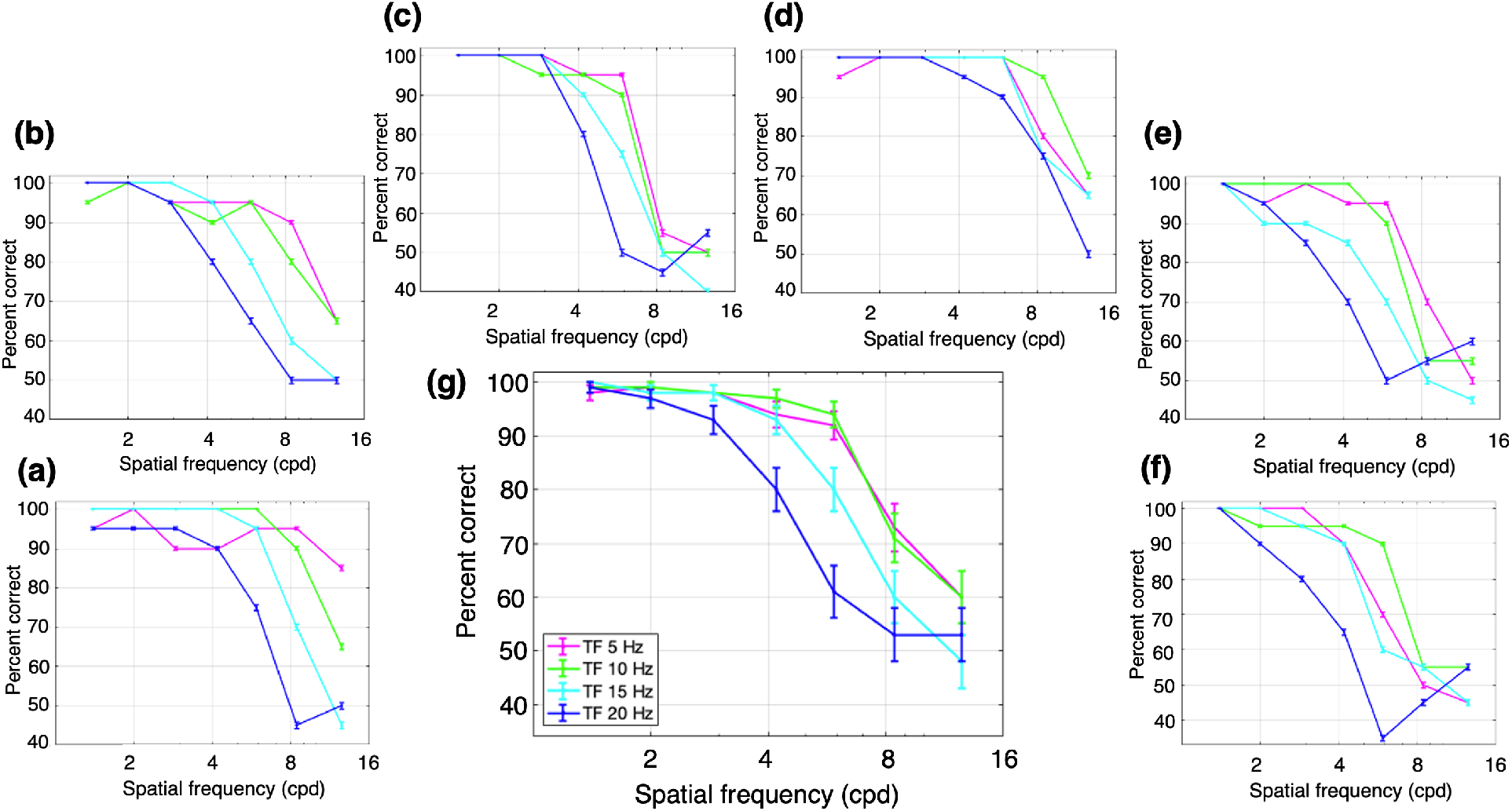
Performance for luminance modulation (LM) stimuli (i.e., drifting sinusoidal gratings). The stimuli consisted of luminance gratings at different SFs and TFs. The task was to discriminate the direction of motion of a grating presented monocularly at 4.3° of eccentricity. Percent correct responses were measured while varying SF (cpd) within blocks for different values of TF: magenta, 5 Hz; green, 10 Hz; cyan, 15 Hz; and dark blue, 20 Hz. **(a-f)** Percent correct responses for each subject. **(g)** Mean percent correct responses of the six subjects. Error bars represent the binomial standard error (SE) of each condition for each subject **(a-f)** and the SEM (N=6 subjects) for the subject-averaged results **(g)**.

In general, the performance for LM stimuli at relatively high SFs was best at low TFs, and decreased with higher values of SF or TF. These findings replicate well-known results for LM stimuli as functions of SF and TF in central vision, where performance at high SFs is best at low TFs, and vice versa^25-27^.

Next, we examined performance with the CM stimuli which had a contrast-reversing grating carrier with right-oblique orientation (45°). The carrier contrast (*C*_*c*_ in Eq 3) was modulated by a vertical drifting sinewave grating envelope with a modulation depth (*C*_*e*_ in Eq 4) of 80%, SF of 0.5 cpd, and TF of 3 Hz. The carrier contrast was 80%, chosen based on pilot experiments to provide roughly similar ranges of performance as for the LM stimuli. Percent correct performance to report the direction of envelope motion is plotted as a function of carrier SF for one subject in **Figure 4a**, for a series of carrier TFs. Performance was generally best at high carrier SFs (2.9 to 5.9 cpd) that were higher than the SFs giving best performance for LM stimuli (**Fig. 3**). Another difference from the LM results is that CM performance at higher SFs was more robust with increasing carrier TF. **Figure 4b-f** plots the results for the 5 other subjects, who showed largely similar results. The average performance across all 6 subjects (**Fig. 4g**) shows a bandpass dependence on carrier SF, which is very similar for the different carrier TFs, with the curves almost overlapping. These observations were confirmed with a 2-way ANOVA, which showed a statistically significant effect of carrier SF, F(6,168) = 70.27, p = 2.97×10^−43^, but no significant effect of carrier TF, F(3,168) = 2.02, p = 0.11, or interaction of carrier SF and TF, F(18,168) = 0.76, p = 0.75.

**Figure 4.**
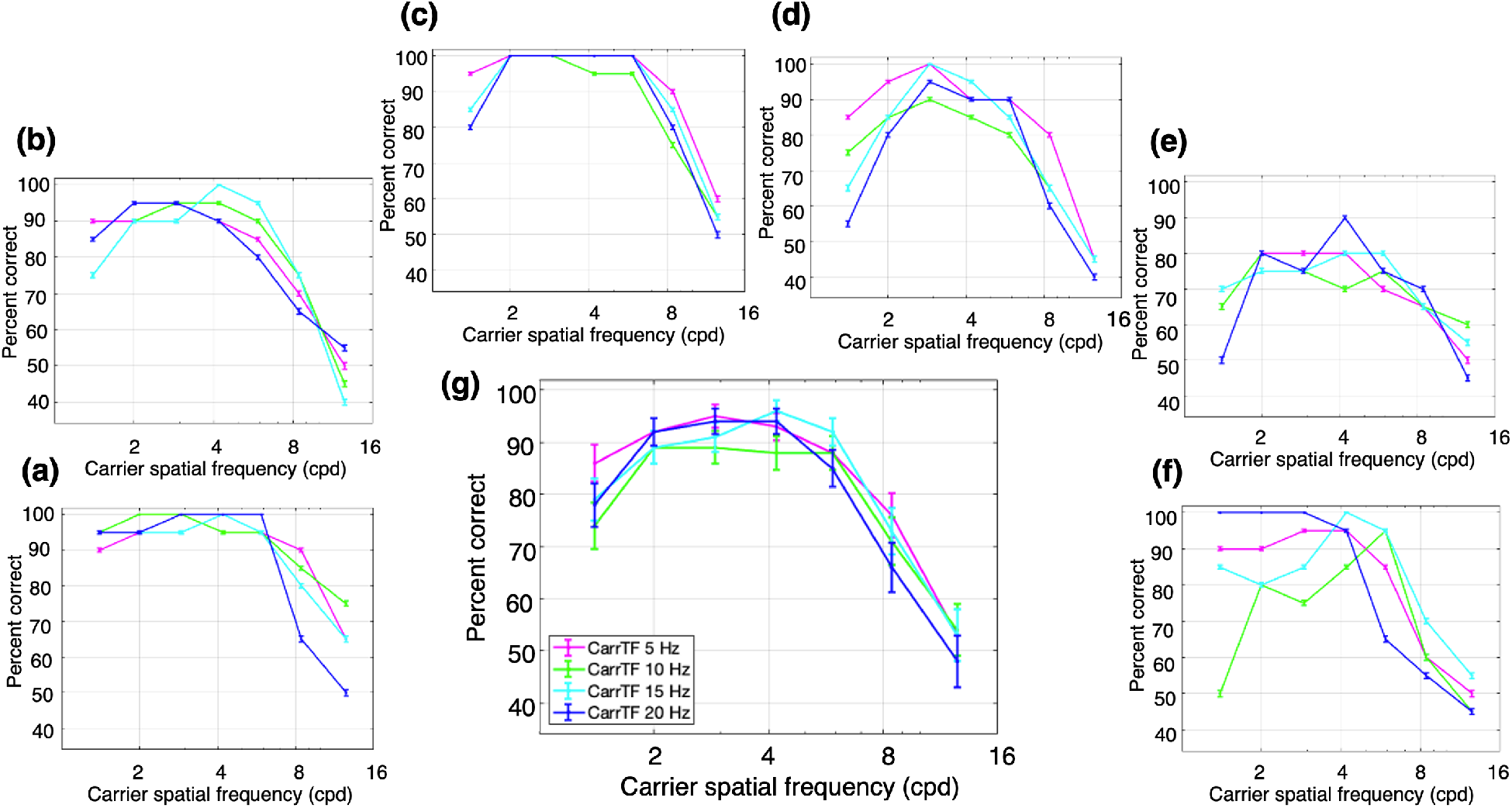
Performance for contrast modulation (CM) stimuli. The stimuli consisted of a contrast-reversing grating (carrier) modulated by a drifting sinewave envelope with a SF of 0.50 cpd and a TF of 3 Hz. The task was to discriminate the direction of motion of the envelope of a CM stimulus presented monocularly at 4.3° of eccentricity. Percent correct responses were measured while varying carrier SF (cpd) within blocks for different values of carrier TF: magenta, 5 Hz; green, 10 Hz; cyan, 15 Hz; and dark blue, 20 Hz. **(a-f)** Percent correct responses of the six subjects. (**g**) Mean percent correct responses of the six subjects. Error bars represent the binomial SE of each condition for each subject (**a-f**) and the SEM (N=6 subjects) for the subject-averaged results **(g)**.

As a whole, the performance for CM stimuli was better at higher carrier SFs and TFs than for LM stimuli. In particular, there was a large degree of independence between the effects of carrier SF and TF. These results would not be expected based on the linear mechanisms that detect LM stimuli, which support good performance when high SFs are combined with low TFs, or vice versa^25-27^. Importantly, the current results for CM stimuli (**Fig. 4**) are in sharp contrast to those for LM stimuli (**Fig. 3**) over the same ranges of spatiotemporal frequencies, consistent with the idea that the processing of CM stimuli is fundamentally different from that of luminance stimuli.

In comparing the results for LM and CM stimuli, differences in the highest SFs giving good performance (i.e., thresholds describing SF acuity) provides a useful way to quantitatively compare LM and CM performance in a concise manner. To this end, we obtained measures of SF acuity (threshold) from the individual subject data in **Figures 3 & 4** by curve-fitting the high-SF fall-off and taking the value of SF corresponding to 75% correct as an estimate of SF acuity (see **Materials and Methods**). Thresholds were successfully obtained for 23/24 and 22/24 of the individual subject curves in **Figures 3 & 4** respectively. The results of this analysis show that SF acuity for LM stimuli (orange curve) declines systematically with TF whereas carrier SF acuity for CM stimuli (purple curve) remains relatively constant with carrier TF (**Fig. 5**). To quantify this assessment, we regressed the log SF acuity on the TF and stimulus type (LM or CM) as well as their interaction. A significant main effect of TF was observed, F (1,41) = 9.45, p = 3.7 ×10^−3^, as well as a significant interaction between TF and stimulus type, F(1,41) = 5.85, p = 0.02, such that SF acuity declined faster with increasing TF for LM stimuli (b = -0.04) than CM stimuli (b = -0.005).

**Figure 5.**
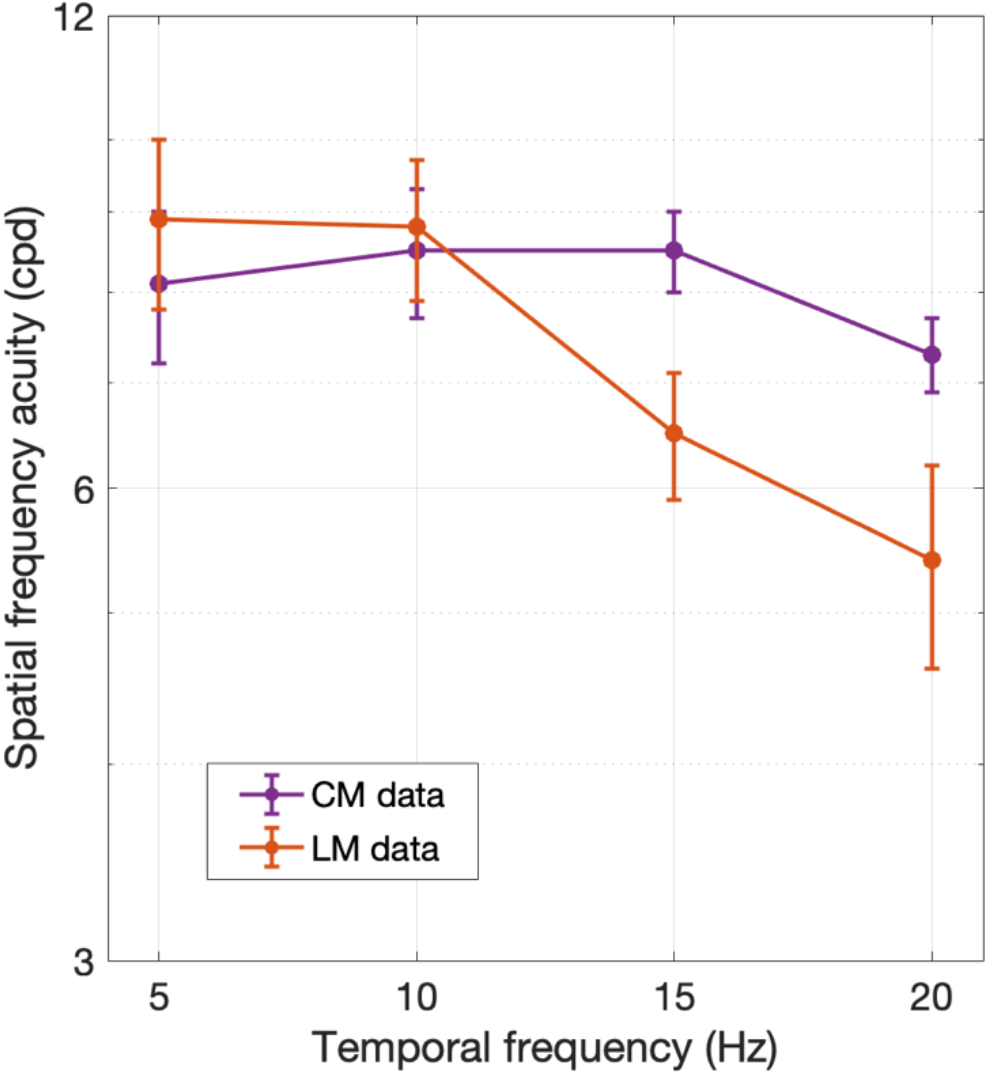
Spatial frequency acuity (i.e., SF corresponding to 75% correct) obtained by fitting the high-SF fall-off of the individual subject data in **Figures 3** and **4** for LM (orange) and CM (purple) stimuli, respectively. Error bars indicate ± SEM (N=6 subjects).

### Experiment 2: Spatial frequency dependence at different eccentricities

Based on previous retinal neurophysiology^8^, we further hypothesized that direction discrimination performance for CM stimuli would remain relatively constant as a function of stimulus eccentricity if the envelope motion was computed from the output of nonlinear Y-like cells rather than linear mechanisms. To test this prediction, we performed a second experiment in which direction discrimination performance for CM stimuli was measured as a function of carrier SF, with a fixed carrier TF, at different eccentricities. Similar to Experiment 1, we compared the results to those with LM stimuli over the same values of SFs, TFs, and retinal eccentricities. For LM stimuli, previous findings would predict a systematic fall-off of performance for higher SFs at increasing eccentricities^28,29^.

In each stimulus presentation of 250 ms duration, an LM or CM stimulus was presented in the center of the screen while the subject fixated a target that was 2.1°, 4.3°, 6.4°, or 8.5° away from the stimulus (**Fig. 2a,d**). The task was to report the direction of motion of the LM grating or of the CM envelope. Different eccentricities were tested in separate trial blocks. Within each block, the SF of the LM grating or the carrier SF of the CM stimulus was varied using the method of constant stimuli with a fixed TF (for LM stimuli) or carrier TF (for CM stimuli) of 20 Hz, respectively. This high value of TF or carrier TF was specifically chosen to minimize the contribution of linear (luminance-based) mechanisms, based on the results of the previous section. Furthermore, for the CM stimuli, the envelope TF was fixed at 3 Hz. This low value was chosen based on the envelope TF tuning of LGN Y-cells which closely match the TF tuning for contrast-reversing gratings^14^ that are widely used to characterize Y-like nonlinearities^7,8,13^. In stark contrast to their low envelope TF preferences, Y-cells often respond best to high carrier TFs (at least as high as 20 Hz), and the preferences for the two TFs are not significantly correlated^15^.

As in Experiment 1, the LM stimuli (vertical gratings, drifting leftwards or rightwards) were presented with a Michelson contrast of 5%. Percent correct performance to report the direction of motion of the grating as a function of SF is plotted for one subject in **Figure 6a** for each of a series of eccentricities. Note that the performance declined with SF, and the high-SF fall-off decreased with stimulus eccentricity. **Figure 6b-f** plots the results for the 5 other subjects, who showed largely similar results. The average performance across all 6 subjects is plotted in **Figure 6g**, where a systematic fall-off in performance at successively higher SFs and eccentricities is evident. These interpretations were confirmed with a 2-way ANOVA (**Fig. 6g**), showing a significant main effect for SF, F(6,168) = 132.67, p = 4.58×10^−61^, a significant main effect for eccentricity F(3,168) = 15.16, p = 8.88×10^−9^, and a significant interaction effect F(18,168) = 2.28, p = 0.003.

**Figure 6.**
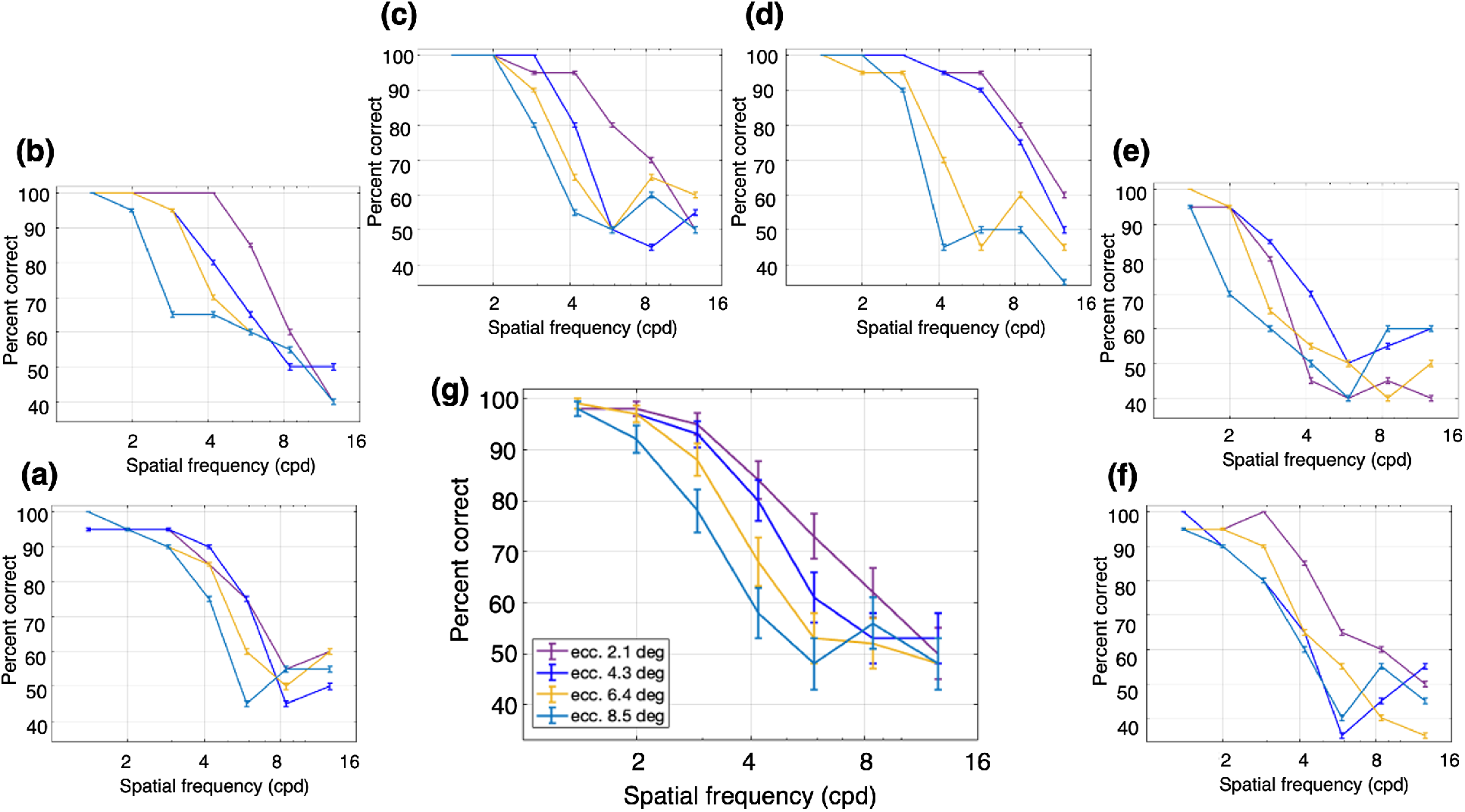
Performance for luminance modulation (LM) stimuli at different retinal eccentricities and a fixed TF of 20 Hz. The stimuli consisted of a luminance grating at different SFs and a fixed TF. The task was to discriminate the direction of motion of the grating when presented at different retinal eccentricities. Percent correct responses were measured while varying SF within blocks for different eccentricities: dark purple, fixation at 2.1°; dark blue, 4.3°; yellow, 6.4°; and light blue, 8.5°. **(a-f)** Percent correct responses for each subject. **(g)** Mean percent correct responses of the six subjects. Error bars represent the binomial SE of each condition for each subject **(a-f)** and the SEM (N=6 subjects) for the subject-averaged results (**g**).

Across subjects the performance for LM stimuli decreased systematically with higher SF and greater eccentricity. These results are consistent with what is expected for LM stimuli, where the dependence of contrast sensitivity as a function of SF shifts to lower SFs as eccentricity increases^28,29^.

Following Experiment 1, the CM stimuli had a contrast-reversing grating carrier (20 Hz) with right-oblique orientation (45°) and a carrier contrast of 80%. The carrier contrast was modulated by a vertical drifting sinewave grating envelope with a modulation depth of 80%, SF of 0.50 cpd, and TF of 3 Hz. Percent correct performance to report the direction of envelope motion as a function of carrier SF at a fixed carrier TF (20 Hz) is plotted for one subject in **Figure 7a** for each of a series of eccentricities. The best performance was at mid-range carrier SFs (2.9 to 5.9 cpd) and was relatively independent of eccentricity compared to the results for LM stimuli (**Fig. 6**). **Figure 7b-f** plot the results for the 5 other subjects who generally showed a bandpass dependence on carrier SF. The average performance across all 6 subjects (**Fig. 7g**) shows this bandpass dependence on carrier SF, with the curves for the different eccentricities so similar as to be largely overlapping. These impressions were confirmed with a 2-way ANOVA (**Fig. 7g**), which showed a significant main effect for the carrier SF, F(6,168) = 55.08, p = 3.43×10^−37^, but no significant main effect for eccentricity F(3,168) = 0.84, p = 0.48, and no significant interaction effect F(18,168) = 0.92, p = 0.56.

**Figure 7.**
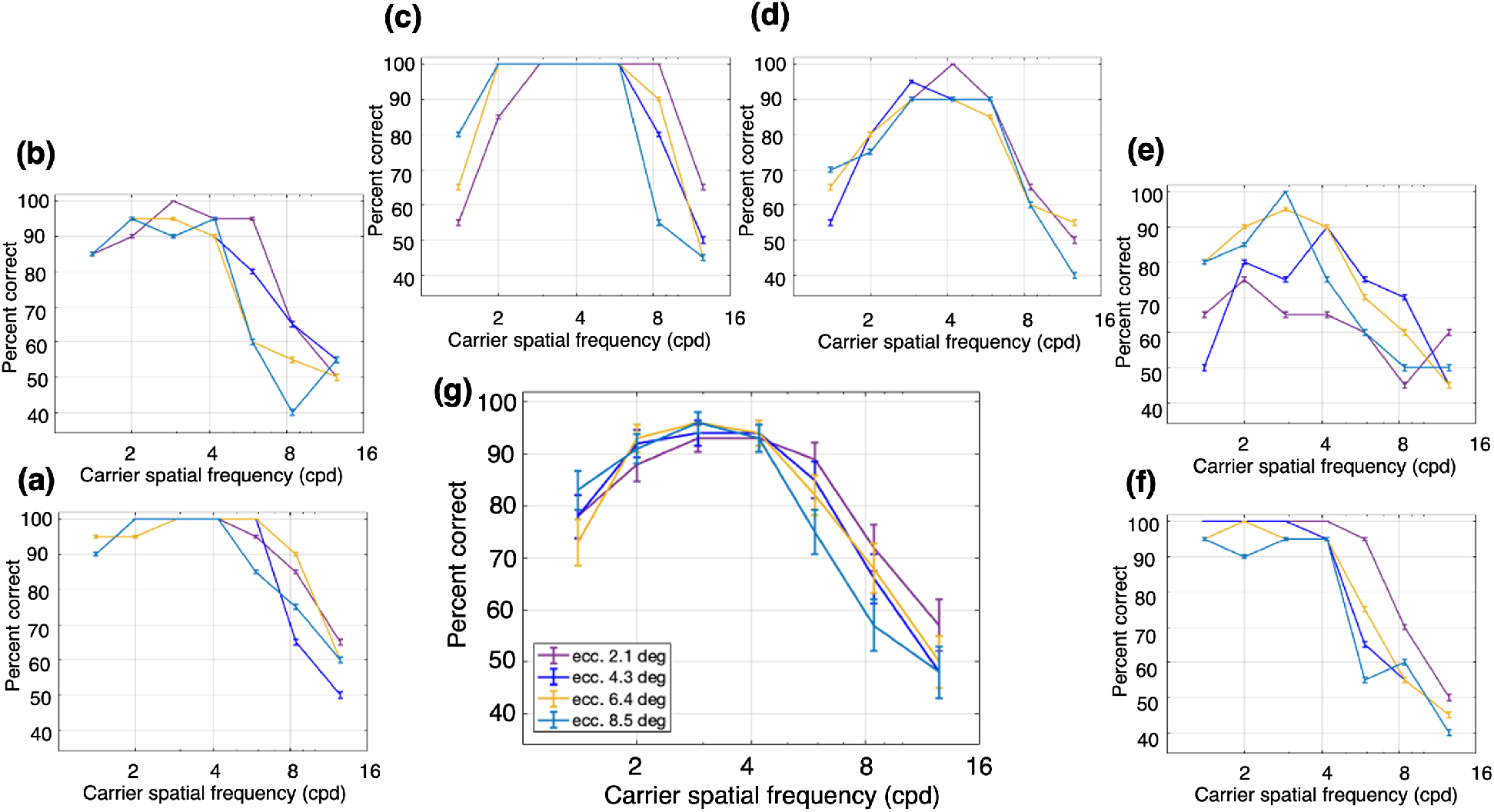
Performance for contrast modulation (CM) stimuli at different retinal eccentricities and a fixed carrier TF of 20 Hz. The stimuli consisted of a contrast-reversing grating (carrier) modulated by a drifting sinewave envelope with a SF of 0.5 cpd and a TF of 3 Hz. The task was to discriminate the direction of motion of the envelope when presented at different retinal eccentricities. Percent correct responses were measured while varying carrier SF within blocks for different eccentricities: dark purple, fixation at 2.1°; dark blue, 4.3°; yellow, 6.4°; and light blue, 8.5°. (**a-f**) Percent correct responses for each subject. **(g**) Mean percent correct responses of the six subjects. Error bars represent the binomial SE of each condition for each subject (**a-f**) and the SEM (N=6 subjects) for the subject-averaged results (**g**).

Notably, the performance with CM stimuli exhibited relatively little dependence on eccentricity (**Fig. 7**), whereas the results with LM stimuli showed a clear, systematic eccentricity-dependence (**Fig. 6**). Only the latter is expected based on a linear mechanism, since the SF-dependence of contrast sensitivity shifts to lower SFs with increasing eccentricity^28,29^. Thus, these results are consistent with the idea that the processing of CM stimuli is fundamentally different (nonlinear) from that of LM stimuli.

Similar to Experiment 1, a useful way to compare LM and CM performance in a concise manner was to measure the differences in highest SFs giving good performance. Consequently, we obtained measures of SF acuity (threshold) from the individual subject data in **Figures 6 & 7** by curve-fitting the high-SF fall-off and taking the value of SF corresponding to 75% correct as an estimate of SF acuity (see **Materials and Methods**). Thresholds were successfully obtained for 18/24 and 23/24 of the individual subject curves in **Figures 6 & 7** respectively. The results of this analysis show that SF acuity for LM stimuli (orange) declines systematically and substantially with eccentricity, whereas the carrier SF acuity for CM stimuli (purple) remains relatively constant, or declines very little, with eccentricity (**Fig. 8**). To quantify this assessment, we regressed the log SF acuity on the eccentricity and stimulus type (LM or CM), as well as their interaction. Consistent with our observations, the effect of eccentricity on SF acuity was significant, F(1,37) = 29.57, p = 3.63×10^−6^, and was twice as large for LM stimuli (b = -0.10) compared to CM stimuli (b = -0.05), although the interaction did not reach statistical significance, F(1,37) = 2.91, p = 0.10.

**Figure 8.**
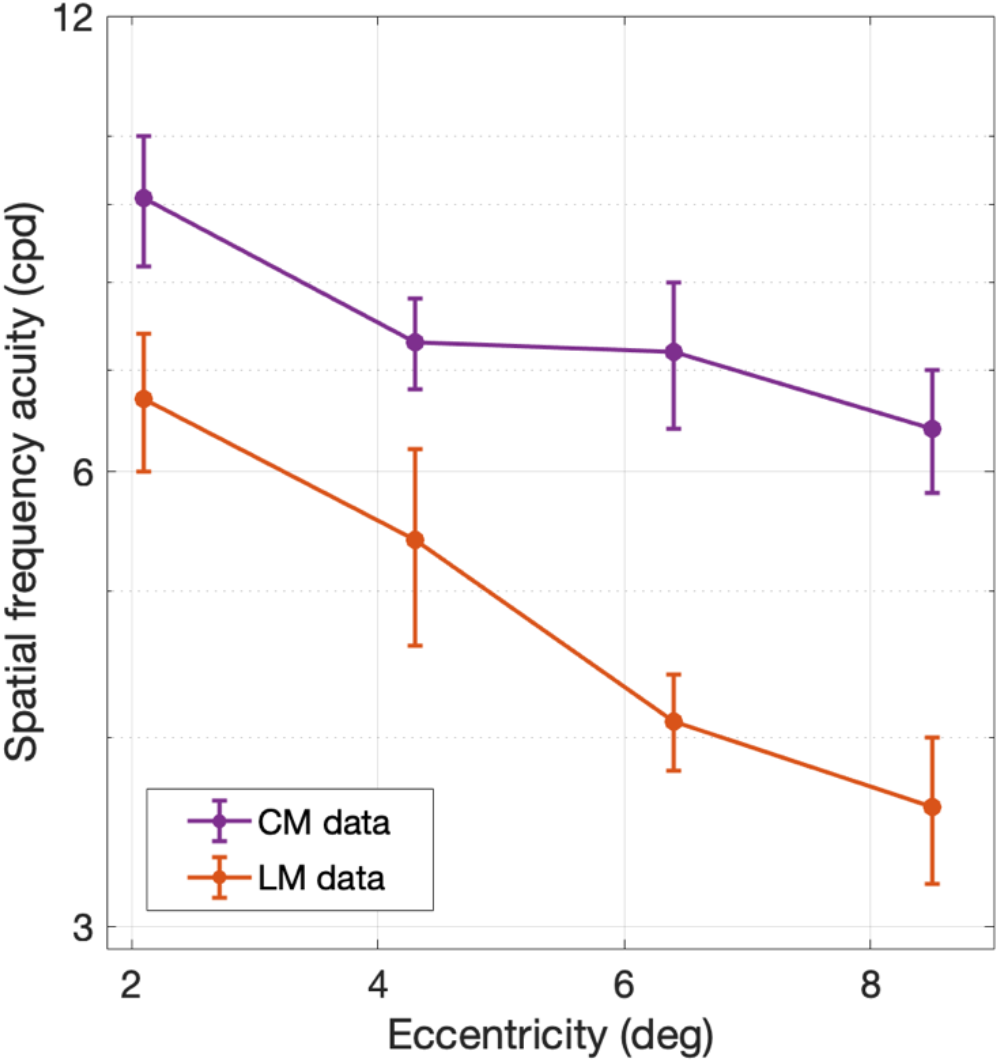
Spatial frequency acuity (i.e., SF corresponding to 75% correct) obtained by curve-fitting the high-SF fall-off of the individual subject data in **Figures 6** and **7** for LM (orange) and CM (purple) stimuli, respectively. Error bars indicate ± SEM (N=6 subjects). Data points for CM stimuli are shifted slightly relative to the actual eccentricity values for visualization purposes.

## Discussion

The encoding of “second-order” stimuli, such as the CM patterns used here, are of particular interest in the context of neural mechanisms underlying perception, since figure-ground boundaries in natural scenes often entail modulations in texture properties such as contrast^30^. Here we introduced a way to assess the subcortical inputs to CM processing in human perception, which was previously possible only in animal models.

CM stimuli have been found to drive early visual cortex neurons in cats^31^ and macaque monkeys^32^, with selectivity to the carrier spatial frequency. The responses to CM stimuli in both cat LGN^14^ and area 18^15,33^ occur at surprisingly high carrier TFs, suggesting a subcortical origin, likely retinal Y-cells. Using CM stimuli at high carrier spatiotemporal frequencies, here we translated this evidence from previous neurophysiology experiments into a human behavioral approach. The results of these experiments suggest that human motion perception of CM stimuli is mediated by carrier processing with nonlinear subunits of Y-like subcortical cells^14,15^, rather than by conventional linear mechanisms thought to detect LM stimuli.

Y-like cells, such as retinal parasol cells, are characterized by their ability to respond linearly to low SF drifting gratings and nonlinearly to contrast-reversing gratings at high spatiotemporal frequencies (e.g., Hochstein & Shapley^4^; Crook et al.^7^). To drive linear mechanism responses, here we employed drifting LM stimuli at different SFs and TFs, which are known to elicit the best behavioral response at combinations of low TF and high SF, or vice versa^25-27^. Not surprisingly, the performance that we found for LM stimuli was consistent with this idea (**Fig. 3**). However, CM stimuli are thought to be specifically processed by nonlinear mechanisms that arise from retinal bipolar cell inputs to the Y-like retinal ganglion cells that project to the LGN^16,34^, whose responses are then relayed to the visual cortex. The nonlinear subunits of LGN Y-cells have been demonstrated to respond to CM stimuli whose individual grating components have higher spatiotemporal frequencies than those that drive linear cortical mechanisms^14,15^. Therefore, nonlinear (carrier) responses of cortical neurons to CM stimuli should be able to produce good behavioral performance at surprisingly high carrier spatiotemporal frequencies. This prediction is consistent with the results in our experiments with CM stimuli, where performance was best at relatively high carrier SFs and high carrier TFs (**Fig. 4**). In addition, the demonstration of a bandpass dependence on carrier SF (**Fig. 4**) is consistent with similar findings for CM-responsive single neurons in early visual cortex of cats^14,31,35^ and macaque monkeys^32^, and with the bandpass SF dependence of nonlinear subunit responses of parasol retinal ganglion cells to simple contrast-reversing gratings^7,8^. The relatively good psychophysical performance for CM stimuli at high carrier TFs (**Fig. 4**) is also consistent with the neurophysiological responses of LGN Y-cells and cortical CM-responsive neurons in the cat^14,15^. While carrier TF tuning properties have not been examined in the primate LGN or visual cortex, the striking similarities of Y-like cells for other stimulus properties across mammalian species^5^ suggest that they will be similar.

Previous human behavioral studies using luminance gratings demonstrated that the SF-dependence on contrast sensitivity shifts to lower SFs with increasing eccentricity^28,29^, much as we found in our experiments with LM stimuli (**Fig. 6**). However, for CM stimuli, we found that performance as a function of carrier SF changed relatively little with eccentricity. This relative invariance of carrier SF acuity to eccentricity might reflect the organization of the nonlinear (F2, second harmonic) receptive fields of Y-like cells. Indeed, previous primate retinal neurophysiology found that the estimated sizes of the center F2 receptive fields of Y-like cells varied minimally with eccentricity, while estimates of the size of the linear (F1, first harmonic) center receptive fields decreased substantially with eccentricity^8^.

Notably, in the experiments of Crook et al.^8^, the eccentricities in the primate retina used to measure F1 and F2 center receptive fields (7°-40°; see their Fig. 6B) were mostly higher than the eccentricities (ca 2°-8°) examined here. While it can be challenging for inexperienced subjects to perform perceptual judgments in more peripheral vision, it nevertheless might be worthwhile in future studies to test a higher range of eccentricities, to make further comparisons to the results of Crook et al.^8^. Based on our results here, and our conclusion that behavioral responses to CM stimuli are mediated by the nonlinear subunits of Y-like cells, we would expect performance to continue declining at larger retinal eccentricities for the LM stimuli at a greater rate than for the CM stimuli.

One can conceivably argue that for our results with CM stimuli, the bandpass carrier SF dependence, in particular the fall-off at low carrier SFs (**Figs. 4, 7**), might be secondary to a lack of a constant ratio between the carrier and envelope SFs, because the envelope SF was kept at a fixed value. In this view, if the carrier and envelope SFs become more similar, it might interfere with the ability to judge the direction of envelope motion as distinct from the carrier flicker, or even that there might be insufficient cycles of the carrier within each envelope cycle to provide a genuine CM stimulus. However, if we had kept a constant ratio, at the lowest eccentricity the visual stimulus would be so large that part of the stimulus would overlap the fovea and in peripheral vision it would be too small to make a reasonable judgement of envelope motion.

Although in our experiments the ratio was not constant, it was always at least 2.8:1 (at the lowest carrier SF), which would provide a reasonable number of carrier cycles within each cycle of the envelope to enable judgement of the direction of envelope motion.

A possible future research direction might be to employ a task in which subjects identify the orientation of the envelope rather than its direction of motion. Because cortical neurons that encode CM stimuli are selective for the orientation as well as the direction of motion of the envelope^32^, this approach should be similarly specific for downstream, cortical processing of Y-like input.

A natural concern when using CM stimuli is that an early nonlinearity in the display screen, or in the photoreceptors, might act to provide an artifactual luminance signal. We performed a careful measurement of the CRT display nonlinearity and incorporated that into the gamma correction of the video attenuator. Then we used the photometer to verify the linearity of luminance with intensity of the corrected video signal. The visual stimuli were always displayed in the center of the CRT (where calibration for gamma correction was performed), with eccentricity being varied by placement of the fixation target rather than the stimulus. After gamma correction, we also employed a diffusing sheet that acts as a spatial low-pass filter to verify that no luminance artifact was visible, thus indicating the calibration was successful in preventing nonlinear distortions^36^. To prevent possible luminance artifacts from the “adjacent pixel nonlinearity"^23,37^, we used sinusoidal carrier stimuli that changed luminance smoothly along the line scan of the CRT display. Furthermore, if the responses to CM stimuli were due to either CRT or photoreceptor nonlinearities, one would not expect a bandpass dependence on carrier spatial frequency (**Figs. 4, 7**). In particular, the fall-off at low carrier SFs demonstrates that the mechanism underlying the CM responses is not secondary to simple luminance artifacts.

The results of this study suggest that tasks employing CM stimuli with high spatiotemporal carrier frequencies can reveal the nonlinear contributions of retinal Y-like cells to human perception. With this approach, we can reveal the SF-selective nonlinear behavior of Y-like cells, separate from the contributions to visual processing from other RGCs. To our knowledge, this is the first human study to employ CM stimuli at different eccentricities with the aim of isolating the nonlinear response function of Y-like RGCs. Based on these results, it is conceivable that the current stimuli may have diagnostic value for disorders which have been linked to deficits in magnocellular function, such as glaucoma^38,39^ and developmental dyslexia^40,41^.

Previous studies suggest that “first-order” stimuli, which are based on variations in luminance, and “second-order” stimuli, which are based on differences in local contrast and texture (such as a CM stimulus)^42^, engage distinct processing mechanisms^23,42,43^. Our results suggest that CM processing likely arises from Y-like cells, and is consistent with neurophysiological evidence from the responses of neurons in the LGN and early visual cortex to CM stimuli^13-15^. Since Y-like cells carry a mixture of first- and second-order information, an important future research direction will be to explore the relationship between the mechanisms underlying the results here and the various other behavioral findings that have supported either common or separate processing for first- and second-order stimuli^36,44-46^.

## Acknowledgments

We thank Chang’an Zhan and Liang Zhen, who each provided invaluable help with technical issues. This work was supported by a Canadian NSERC grant OPG0001978 to CLB and a National Institutes of Health Grant (EY029438) to AR.

## Author contributions

CLB, AR designed the experiment and received funding. ALR collected the data. ALR, LT analyzed and interpreted the data. ALR, CLB, AR wrote and edited the main manuscript. All authors reviewed the manuscript.

## Data availability

The data generated and analyzed for the current study are provided in Supplementary Information file S2.

## Additional information

Correspondence and requests for materials should be addressed to CLB.

